# High-resolution comparative atomic structures of two Giardiavirus prototypes infecting *G. duodenalis* parasite

**DOI:** 10.1101/2023.11.01.564773

**Authors:** Han Wang, Marucci Gianluca, Anna Munke, Mohammad Maruf Hassan, Marco Lalle, Kenta Okamoto

**Affiliations:** The Laboratory of Molecular Biophysics, Department of Cell and Molecular Biology, Uppsala University, Uppsala, Sweden; Unit of Foodborne and Neglected Parasitic Diseases, Department of Infectious Diseases, Instituto Superiore di Sanità (ISS), Rome, Italy; Center for Free-Electron Laser Science CFEL, Deutsches Elektronen-Synchrotron DESY, Notkestr.85, 22607 Hamburg, Germany

**Keywords:** G. duodenalis, dsRNA virus(es), capsid structure, cryo-electron microscopy (cryo-EM), Totiviridae, Giardia lambia virus (GLV)

## Abstract

*Giardia lamblia virus* (GLV) is a non-enveloped icosahedral dsRNA virus and an endosymbiont virus infecting the zoonotic protozoan parasite *Giardia duodenalis* (syn. *G. lamblia, G. intestinalis*), a pathogen of mammals, including humans. Elucidating the transmission mechanism of GLV is crucial to an in-depth understanding of the virulence of the virus in *G. duodenalis*. GLV belongs to the family *Totiviridae,* which infects yeast and protozoa intracellularly; however, it also transmits extracellularly, similar to phylogenetically distantly related toti-like viruses that infect multicellular hosts. The GLV capsid structure is extensively involved in the longstanding discussion concerning the acquisition of extracellular transmission in *Totiviridae* and toti-like viruses. Hence, this study constructed the first high-resolution comparative atomic models of two GLV strains, namely GLV-HP and GLV-CAT, which showed different intracellular localization and virulence phenotypes, using cryo-EM single-particle analysis. The atomic models of the GLV capsids presented swapped C-terminal extensions, extra surface loops, and a lack of cap-snatching pockets, similar to those of toti-like viruses. However, their open pores and lack of the extra crown protein (CrP) resemble those of other yeast and protozoan *Totiviridae* viruses, demonstrating the essential structures for acquiring extracellular cell-to-cell transmission. The intensive structural comparison between GLV-HP and GLV-CAT indicates the first evidence of critical structural motifs for the transmission and virulence of GLV in *G. duodenalis*.

## Introduction

*Giardia duodenalis* (syn. *G. lamblia* and *G. intestinalis*) is a zoonotic intestinal protozoan parasite that infects the upper part of the small intestine of mammals, including humans. *G. duodenalis* causes giardiasis, a widespread diarrheal disease in humans [1]. The pathogen is classified into eight distinct genetic groups or assemblages (A–H), with human infection almost exclusively associated with Assemblages A and B, whereas giardiasis in animals is due to host-specific Assemblages (C–H). Assemblages A and B also have zoonotic potential, having been isolated from both humans and animals infected with *G. duodenalis* [2]. Transmission of *G. duodenalis* occurs through the fecal-oral route by accidental ingestion of cysts, the infective environmental resistant stage of the parasite, by direct contact with stools of humans and animals infected with the parasite, or by drinking water or eating food (e.g., fresh produce) contaminated with the parasite’s cysts. Outbreaks of giardiasis are frequently reported globally [3,4] as well as recently in EU countries in 2018–2019 [5,6]. However, infection with *Giardia* is still considered a neglected disease; thus, effective anti-parasitic drugs are limited [7], vaccines are not yet available [8,9], and treatment failure with nitroimidazoles is increasingly reported, with up to 45% of patients not responding to initial treatment [10].

For neglected human protozoan parasitic infections, virotherapy using parasite-specific endosymbiont viruses has been proposed as an alternative approach to controlling them [11–13]. The double-stranded (ds)RNA virus Giardia lamblia virus (GLV or *Giardiavirus*) is able to infect *G. duodenalis* [14,15]. GLV belongs to the genus Giardiavirus in the family *Totiviridae*. Other *Totiviridae* viruses include diverse dsRNA viruses that exclusively infect yeasts and fungi, such as the Saccharomyces cerevisiae virus L-A (ScV-L-A), the Saccharomyces cerevisiae virus L-BC (ScV-L-BC), and the Helminthosporium victoriae virus 190S (HvV190S), or that infect protists such as Trichomonas vaginalis virus (TVV) (genus Trichomonasvirus) and Leishmania spp. RNA virus (LRV) (genus Leishmanivirus) [15–17]. The genetically distantly related toti-like viruses, for example, mosquito Omono River virus (OmRV), shrimp infectious myonecrosis virus (IMNV), and salmon piscine myocarditis virus (PMCV), infect a broad range of invertebrates and vertebrates [16,18,19].

GLV is phylogenetically close to toti-like viruses compared to other *Totivirdiae* viruses, which hints at an understanding of the evolutionary relevance of *Totiviridae* viruses and toti-like viruses [16,20]. Almost all *Totiviridae* viruses infecting unicellular yeast and protists merely adopt intracellular transmission by frequent mating of cell division and cell fusion in yeasts and protists [17,21,22], while the toti-like viruses infecting multicellular hosts have acquired an extracellular phase and likely transmit adjacent cells using membrane-penetration and/or receptor-binding mechanisms [23–26]. Intriguingly, although GLV infects a unicellular protozoa, it can be efficiently transmitted extracellularly [15,27].

*Totiviridae* and toti-like viruses encode two open reading frames (ORFs), ORF1 and ORF2, in their genome [15,16]. ORF1 expresses a capsid protein (CP) that assembles an icosahedral capsid. Only for the toti-like viruses is the CP further post-translationally processed to form major capsid protein (MCP) and crown protein (CrP) [16,24]. These viral CP (or MCP) and CrP are in charge of the infection of their hosts [24,28]; however, their role in infection is still poorly understood. The CP of *Totiviridae* viruses has another function: snatching the cap RNA structure of the host. The attachment of the 5’-cap structure (7-methylguanocine linked through reverse 5’-5’ triphosphate bridge; 5’-m^7^GpppX) to mRNA is required for effective translation in eukaryotic cells. Many viruses express enzymes, such as RNA triphosphatase, guanylyltransferase, and methyltransferase, to process viral RNA via several catalytic pathways to create the 5’-cap structure; however, some viruses, especially negative sense single-stranded (-)ss)RNA viruses, take a cap-snatching approach and utilize a short cleaved capped RNA fragment from the host mRNAs [29,30]. The invariant His residue of the CP in the yeast *Totiviridae* viruses binds covalently to the m^7^GpppG cap structure of the host mRNA, thereby decapping the m^7^Gp moiety, which speculates a unique cap-snatching approach of transferring the m^7^Gp to the 5′ end of the synthesized viral positive sense (+)ssRNA transcripts [31,32]. However, for GLV, the transcripts lack the 5’-cap structure, and a cap-independent internal ribosomal entry site (IRES) could promote the translation of the structure [15,33].

ORF2 encodes RNA-dependent RNA polymerase (RdRp), which is expressed in a fused CP-RdRp form by ribosomal frameshifting [15,16]. During capsid assembly, one or a few CP-RdRp(s) are incorporated inside the virus particle [34,35]. The internal RdRps function for intraparticle genome synthesis, which synthesizes nascent (+)ssRNAs and the (+)ssRNAs, should exit the pore(s) on the virus capsid [25,36,37]. Intraparticle genome synthesis and the RdRp/pore are thought to be fundamental requirements of non-enveloped icosahedral dsRNA viruses for sequestering the host’s innate immune system that could be triggered by viral dsRNAs [25,37]. However, the pores of some icosahedral dsRNA viruses that infect multicellular hosts (e.g., toti-like viruses) are obstructed [23,25,38]. Considering the unique phylogenetic clade of GLV in evolution, the surface and pore structure of GLV are key to deeply elucidating the infection and intraparticle genome synthesis mechanisms in the transmission steps of *Totiviridae* and toti-like viruses.

Recently, two subtypes of GLV, namely GLV-HP and GLV-CAT, have been described in depth [15]. These subtypes have been found to infect the *G. duodenalis* Assemblage A isolate HP-1, of human origin, and CAT1, of cat origin. Despite limited divergence at nucleotide and amino acid levels, GLV-HP and GLV-CAT show different phenotypes when individually infecting the naive *G. duodenalis*Assemblage A isolate WBC6 [15]. GLV-HP has the tendency to form aggregates and to accumulate below the trophozoite plasma membrane and inside the cell body, whereas GLV-HP does not form aggregates, is scattered in the trophozoite, and does not accumulate in the cell body [15]. Additionally, chronic infection of GLV-HP hampers parasite growth more markedly than GLV-CAT, indicating that GLV-HP might have features of a pre-lytic virus [15].

The atomic models of the yeast ScV-LA and L-B [35,39], protozoan TVV2 [40], and toti-like virus OmRV [24,25] have recently been determined; however, the structure of GLV is still limited to an intermediate resolution of 6 Å [20]. Here, we describe the first high-resolution structure and atomic model of both GLV-HP and GLV-CAT virions, demonstrating the conserved and unique structural features of infection and intracellular genome synthesis in GLV and other *Totiviridae*/toti-like viruses. The intensive comparison between GLV-HP and GLV-CAT also presents potential structural motifs that could greatly modify the lifestyle and virulence of the GLV strains.

## Results and discussion

### Cryo-EM structural determination, capsid geometry, and atomic models

The first high-resolution capsid structure of two GLV subtypes was determined using cryo-EM single-particle analysis at a resolution of 2.1 Å and 2.6 Å for GLV-HP and GLV-CAT, respectively (Figs. 1A and 1H, Supplementary Fig. S1 and Table S1). As reported previously [15], the GLV-HP particles tend to interact with each other, but the GLV-CAT particles do not (Figs. 1B and 1I). The achieved resolution of the obtained cryo-EM maps is the highest in the *Totiviridae* virus database to date, which enables us to more precisely discuss the capsid structure and CP–CP interactions.

**Fig. 1.**
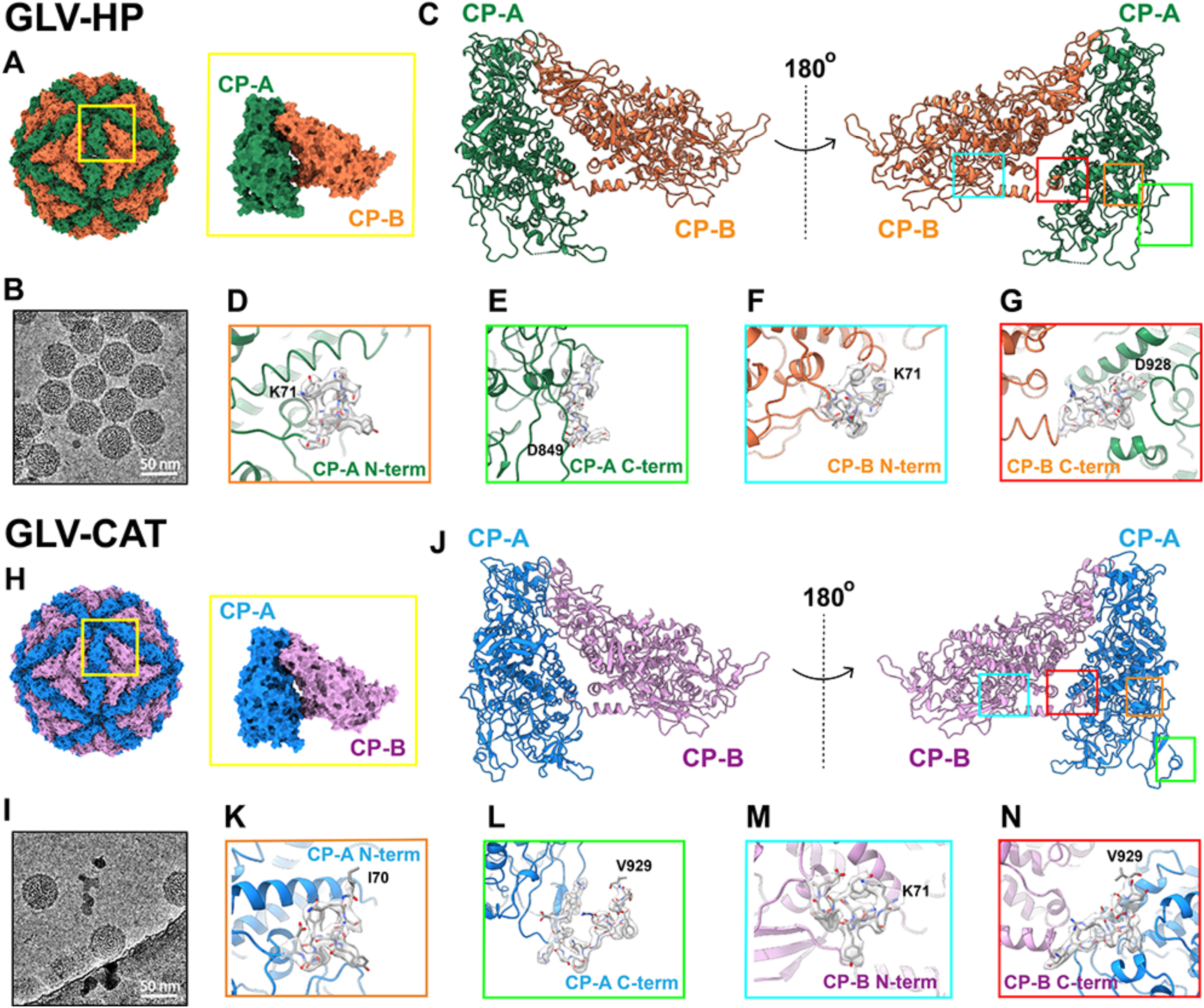
Atomic model of capsid and CPs in GLV-HP and GLV-CAT. The T = 1 capsid geometry and CP-A and CP-B organization in (A) GLV-HP and (H) GLV-CAT. Raw cryo-EM micrographs of (B) GLV-HP and (I) GLV-CAT. The closed-up views of N-termini and C-termini in CP-A and CP-B are shown in (C–G) for GLV-HP and (J–N) for GLV-CAT.

The GLV capsid is composed of 60 copies of CP dimers named A and B subunits (CP-A and CP-B) in the icosahedral asymmetric unit (Figs. 1A, 1C, 1H, and 1J). Considering the predicted internal IRES sequence in the GLV genome, the translation of ORF1 (CP) does not initiate from the first methionine but from an internal amino acid residue [15,33]. The first amino acid residue of the CP was settled from the internal Pro residue (PENIT …), according to a previous mass spectrometry analysis of the purified GLV particles (Supplementary Fig. S2) [15]. The atomic models of CP-A and CP-B, both in GLV-HP and GLV-CAT, were built from the Ile70 or Lys71 residue to Asp928 or Val929 residue, apart from the CP-A of GLV-HP (Figs. 1D-1G, 1K-1N, Supplementary Fig. S2). The N-terminal residues (residues 1–69) of the CP seem to be located on the interior side of the capsid and are not found in the cryo-EM maps (Supplementary Fig. S2). The overall structure of the GLV CP shows typical α-helix-rich α+β fold commonly adopted in the *Totiviridae* and other icosahedral dsRNA viruses (Figs. 1C and 1J, Supplementary Fig. S2). The structures of CP-A and CP-B are very similar, as the total root mean square deviation (RMSD) is 4.338 Å and 8.863 Å in the GLV-HP and the GLV-CAT capsid, respectively (Supplementary Fig. S3); however, the interfaces between the two adjacent subunits make some differences in CP-A and CP-B (red dotted circles in Supplementary Fig. S3).

### C-terminal extension and CP–CP interactions

In the GLV capsid structure, the CP–CP interactions are mainly mediated by the adjacent interfaces of CP-A and CP-B (Fig. 1) and the short C-terminal extension that interlocks CP-A and CP-B on the interior side (Fig. 2). The C-terminus of CP orients differently in CP-A and CP-B due to the interlocking between two adjacent subunits. Hence, this C-terminal interlocking is identified in both directions of CP-A to CP-B and CP-B to CP-A in the GLV-CAT capsid (Figs. 2C–E); however, only the CP-B to CP-A direction is identified in the GLV-HP capsid (Figs. 2A–B). This C-terminal interlocking includes three possible salt bridges: Arg671–Asp922, Arg675–Asp928, and Lys774–Asp927 in CP-B to CP-A in GLV-HP (Fig. 2B). The C-terminal interlocking includes the same amino acid pairs as Arg671–Asp922 and Lys774–Asp927; however, more intersubunit interactions are identified in GLV-CAT (Fig. 2E). In the CP-A to CP-B directions, only two possible salt bridges—Arg675–Asp928 and E872–N917—are present in GLV-CAT (Fig. 2D). In the ScV-L-BC capsid, only a very short C-terminal extension of the CP-A has been identified [35], and ScV-L-A, LRV-1, and TVV2 have no interlocking C-terminal extension [39–41]. Considering the C-terminal structure of these viruses, the C-terminal extension of GLV is longer and interlocks adjacent A and B subunits in both directions, at least in GLV-CAT (Figs. 2C–E). This interlocking mechanism in the GLV capsid is very similar to that of the toti-like virus OmRV. The C-terminal extension of the OmRV AK4 strain swaps the orientation on Ser1603 residue in the A and B subunits to enable interlocking in both the A to B subunits and the B to A subunits [25]. In the OmRV LZ strain, the long C-terminal extension of the B subunit interacts with several neighboring subunits [24]. In the C-terminal arm of the GLV-CAT subunits, similar swapping occurs structurally on the Lys902 residue. Other segmented and large icosahedral dsRNA viruses, the megabirnavirus and the quadrivirus, show an extralong C-terminal extension to interact with adjacent subunits or several proximal subunits [23,42].

**Fig. 2.**
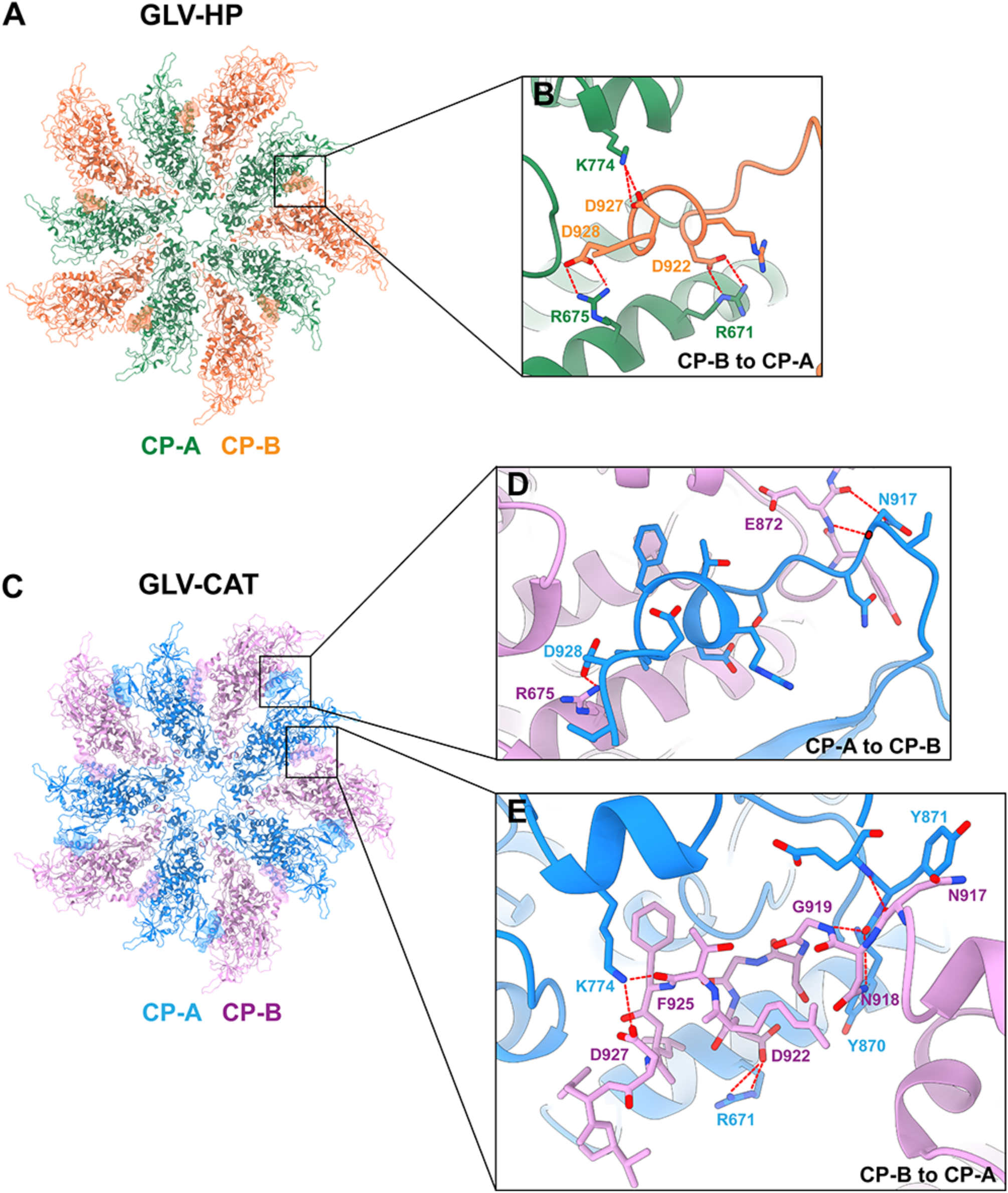
C-terminal extension structure of the CP–CP interface in the GLV-HP and GLV-CAT capsids. The atomic model of the 5-fold CP complex of (A) GLV-HP (CP-A in green; CP-B in orange) and (C) GLV-CAT (CP-A in light blue; CP-B in pink) is shown from their inside view. The interlocking C-terminal extensions from CP-A to CP-B and those from CP-B to CP-A are highlighted in the surface representation. The closed-up views of the interactions between the C-terminal extension and the adjacent capsid are shown in B for GLV-HP and in D and E for GLV-CAT. The amino acid residues involved in the interactions are indicated. Red dashed lines between amino acid residues indicate predicted salt bridges.

It is interesting why the C-terminal extension is required for GLV, OmRV, and the other segmented icosahedral dsRNA viruses. It has been previously hypothesized that the C-term extension may be required for the stability of the larger capsid in the consequence of packaging its own longer genome to tolerate the necessary higher interior pressure [23,25,35]. The capsid of ScV-L-A, ScV-L-BC, LRV-1, and TVV2 is 35–42 nm in diameter and packages a 4.5–5.5 kbp dsRNA [35,39–41], while that of the toti-like virus OmRV is 42 nm in diameter and packages a 7.5 kbp dsRNA [24,25]. The GLV capsid is 45 nm in diameter, packaging 6.2 kbp dsRNA [15], which implies that the interior pressure of the GLV capsid is slightly less than that of the OmRV capsid but more than or similar to those of the ScV-L-A, ScV-L-BC, and TVV2 capsids. Therefore, the length of the GLV C-terminal extension and its contribution to interlocking correspond to the expected interior pressure in *Totiviridae* and toti-like viruses. The C-terminal extension is also important for the interior localization of the RdRp; however, the structure of the fused CP-RdRp has not been determined in the obtained cryo-EM map.

### Structural relations of CP within GLV, *Totiviridae* viruses, and other icosahedral dsRNA viruses

The CPs of T = 1 icosahedral dsRNA viruses share a conserved α-helix-rich α+β structural fold, as does GLV CPs (Fig. 1); thus, their structural alignments can inform unique and common structural features within these viruses. To search for CP structures that are akin to that of GLV in the latest PDB repository, the atomic model of the GLV CP was queried in the Dali online server, yielding eight similar CP structures of dsRNA viruses belonging to *Totiviridae*, including the unclassified toti-like virus OmRV, *Chrysoviridae*, *Megabirnaviridae*, and *Quardriviridae* viruses (Supplementary Fig. S4A). All the obtained atomic models were further aligned using MUSTANG and generated an RMSD-based structure phylogeny of these CPs (Supplementary Fig. S4B). Both results indicate that the closest CP structure of GLV is that of the toti-like virus OmRV and that it is distantly related to those of the other *Totiviridae* viruses infecting yeast and protozoa (Supplementary Fig. S4). Genetic phylogenetic analysis has demonstrated that GLV has its own cluster that is located between *Totiviridae* and toti-like viruses [16,20]. The structural comparison between the CPs of GLV and OmRV CPs provides new insight into the unclarified relationship between *Totiviridae* and toti-like viruses, which might reflect their acquisition of functional structures on the capsid.

Proceeding on this assumption, we intensively compared the CP structure of yeast and protozoan *Totiviridae* viruses with the toti-like virus OmRV. The GLV CP presents several extra surface loops on the capsid surface when compared with those of yeast ScV-L-A and ScV-L-BC and protozoan TVV2, which infect intracellularly (Fig. 3), and the OmRV CP presents similar extra surface loops on the capsid (Fig. 3). These extra surface loops on the OmRV CP have previously been suggested as unique to the toti-like virus CPs and not exhibited in the yeast ScV-L-A CP [25]. These surface loops are located in amino acid residues 156-169, 364-388, 415-443, 527-546, and 565-583 in the GLV CP (red regions in Fig. 3B), and 874-883, 1076-1095, 1115-1141, and 1541-1551 in the OmRV CP (blue regions in Fig. 3B). GLV and OmRV acquired an extracellular phase, unlike other *Totiviridae* viruses, which have a non-exceptional intracellular transmission mechanism [15,16,27].

**Fig. 3.**
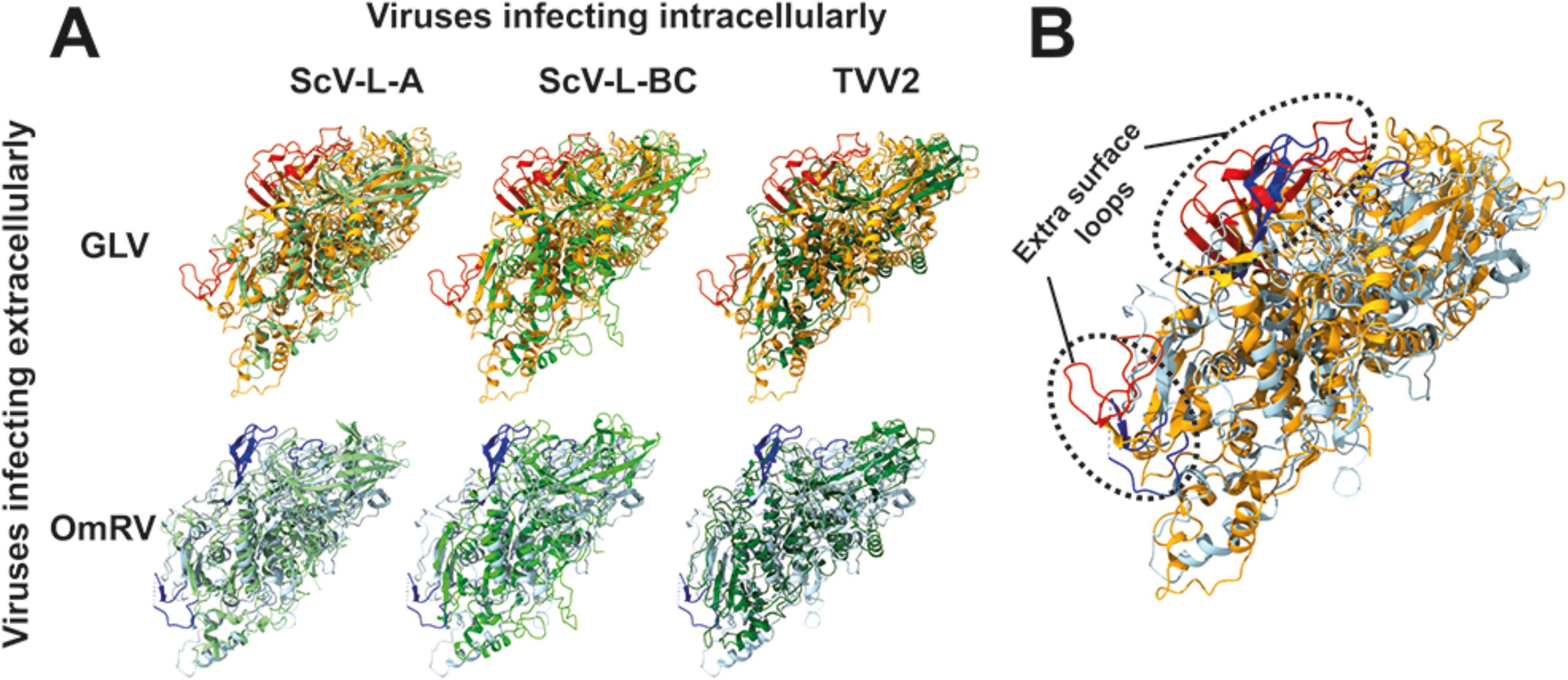
Structural comparison of the GLV CP with those of *Totiviridae* and toti-like viruses. ScV-L-A, ScV-L-BC, and TVV2 CPs are shown in light green, lime, and green. The GLV and OmRV CPs are shown in orange and light blue. The unique surface loops of the GLV CP are shown in red, and those of the OmRV CP are shown in blue. (A) The CP of GLV and OmRV that infect extracellularly is aligned with that of ScV-L-A, ScV-L-BC, and TVV2, which infect intracellularly. (B) The aligned CP structure of GLV and OmRV. The dotted circles indicate the surface regions of the extra loops identified in the GLV and OmRV CPs.

The extracellular transmission mechanism of *Totiviridae* and toti-like viruses is still unclear; however, these surface loops could have profound roles in extracellular cell-to-cell transmission, such as budding, membrane penetration, and receptor-binding cell entry. The toti-like virus OmRV and IMNV and the megabirnavirus RnMBV-1 also possess additional CrP on their surface, which may facilitate extracellular or horizontal cell-to-cell transmission in multicellular hosts [23,24,43]. Considering that the GLV capsid does not express and has no extra surface CrP structurally (Fig. 1), CrP must be a non-essential factor in cell-to-cell transmission. These findings agree with previous results that inhibiting CrP interaction with the OmRV capsid does not completely eliminate the infectivity of OmRV [24,28]. Since the amino acid sequence of CrP is not predicted to be a cell-penetrating peptide, CrP might only be implicated in receptor-binding and subsequent endocytosis. Further, OmRV particles without CrPs are still infectious to host mosquito cells [34]. To fully understand the transmission mechanisms of *Totiviridae* and toti-like viruses, it is necessary to study these identified extra surface loops.

### 5-fold pore structure

The *Totiviridae* viruses have a pore on each 5-fold axis [35,40] or each 2-fold/3-fold axis [44], possibly for exporting cellular nucleotide triphosphates (NTPs) to synthesize and release viral (+)ssRNA in the intrapartically packaged RdRp(s) to cytosol. In *Reoviridae*viruses, the pore on the T = 1 inner capsid dynamically and structurally collaborates with transcribing the nascent viral (+)ssRNAs on the interior RdRp in-situ [36,45–47]. The structural mechanisms of the pores in *Totiviridae* viruses are seldom described due to the lack of their in-situ transcribing structures. This is probably because only a few or one RdRp are randomly incorporated into their capsid particles, unlike *Reoviridae* viruses [35], although a partially spooled genome structure like the *Reoviridae* viruses is seen in capsid-subtracted 2D class projections in our GLV structure (Supplementary Fig. S5).

Both GLV-HP and GLV-CAT capsid structures exhibit pores on each 5-fold axis with an outer and an inner diameter of 10-11 Å and 19-20 Å, respectively (Fig. 4), like other *Totiviridae* viruses [25,35,40]. At the center of the pore, the surface is positively charged because of the cluster of the 10 lysine residues (five Lys209 and five Lys217 residues) (outside view in Fig. 4). The interior side of the 5-fold pore is, on the contrary, negatively charged (inside view in Fig. 4). These positive and negative surface properties are constantly observed in other *Totiviridae* viruses [25,35,40], which implies their essential functions. The dsRNA viruses infecting multicellular hosts possess similar pores composed of two consecutive arginine residues (RR motif); however, they are obstructed [23,25,34,38]. In human picobirnavirus and toti-like viruses, simple conformational changes on the pore assist them in opening [25,38].

**Fig. 4.**
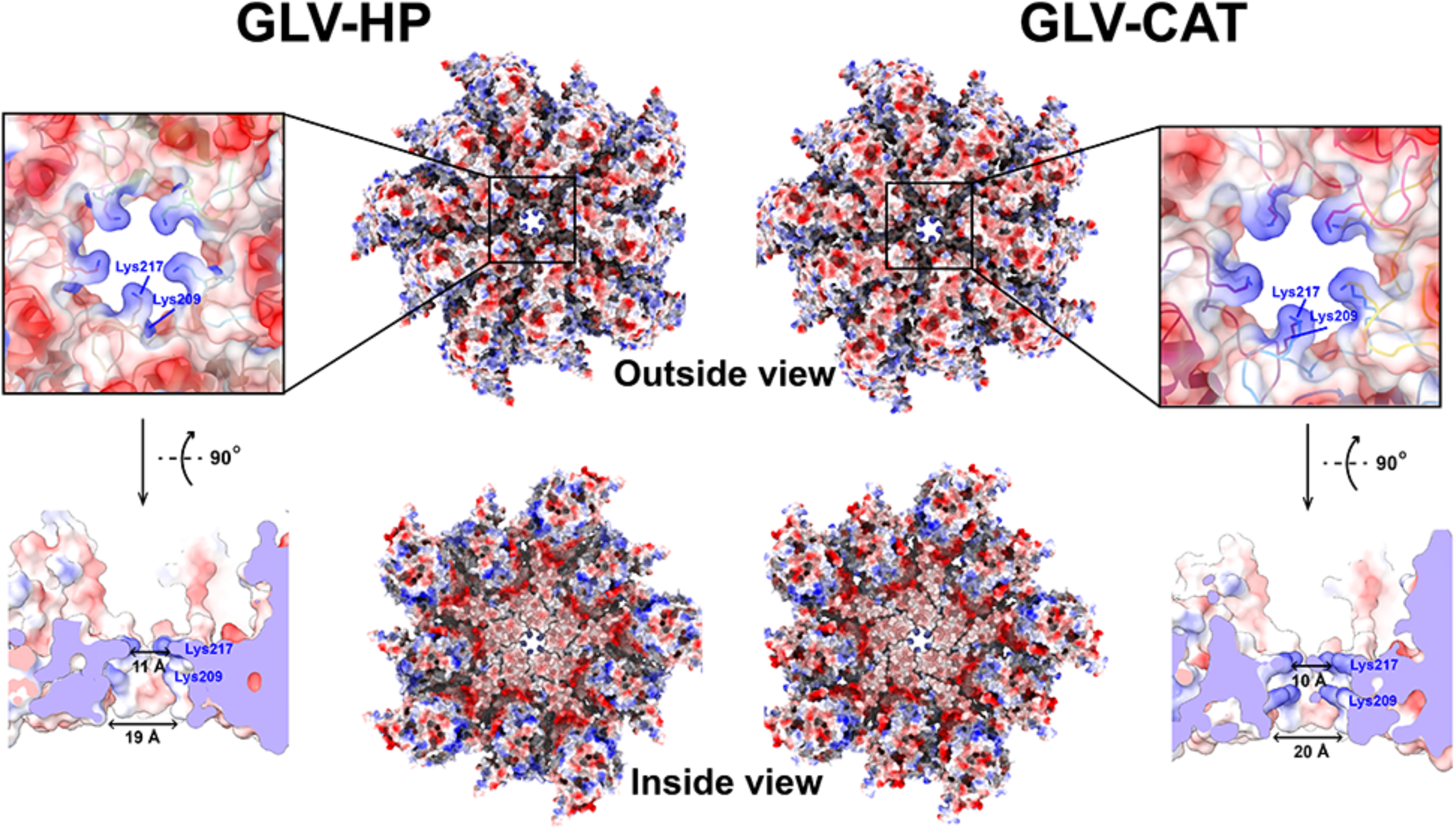
Pore structure of the GLV capsid. A surface electrostatic potential map of the 5-fold CP complex of GLV-HP and GLV-CAT from outside and inside views is shown in red (negatively charged) and blue (positively charged) scales. A close-up view and cross-section of the pore structure and surface charges are also shown at the bottom.

The electropositive capsid surface could have a role in recruiting negatively charged NTPs to the pores, as previously suggested for icosahedral dsRNA viruses [23,25]. A capsid of the human immunodeficiency virus (HIV) also has negatively charged pores for recruiting deoxynucleotide triphosphates (dNTPs) for its intraparticle genome synthesis [48,49]. However, the positively charged residues of the HIV capsid pores are also critical for trapping and utilizing cellular inositol phosphates (IP5 or IP6) during capsid assembly and maturation [50,51]. These findings and previous results imply the multifunctionality and importance of the two lysine residues of the GLV pore.

### Lack of putative cap-snatching active pocket

In yeast *Totiviridae* viruses ScV-L-A and ScV-L-BC, a cap-snatching pocket that exhibits the His residue, His154 in ScV-L-A or His156 in ScV-L-BC, is structurally located on similar regions of the capsid each other [35,39] (Fig. 5A). Although the active pocket is built up with a certain number of amino acid residues, only the invariant His is known to be functional [32]. It is speculated that the protozoan TVV2 CP could have its cap-snatching pocket in a structurally similar location to the yeast CPs [40] (Fig. 5A); however, this is still under discussion. In GLV and OmRV CPs, the cap-snatching pocket and the invariant His residue cannot be found in the position corresponding to that of the yeast ScV-L-A and ScV-L-BC CPs (Fig. 5A). Intensive structural alignments of *Totiviridae* and toti-like viruses revealed that His154 in the ScV-L-A and His156 in the ScV-L-BC CPs are located on the surface region that is followed by structurally well-conserved helices in the *Totiviridae* and toti-like viruses’ CPs (Fig. 5B). Due to the swapped orientation of the secondary structural elements after the structurally conserved helices in TVV2, GLV, and OmRV CPs, their corresponding His residue is not located structurally in the previously reported putative cap-snatching pocket (Fig. 5B). Other viral methyltransferases, such as flavivirus NS5 protein, and cap-snatching complexes, such as influenza virus PA/PB1/PB2 complex, have a cavity formed by aromatic amino acid residues (Trp, Tyr, Phe). These viral enzymes show a negatively charged active site next to the positively charged cluster [29,30,52]. In the cryo-EM map of GLV and OmRV CPs, no such charged surface is obvious in the surface area of the His residues (His320, His419, and His476 in GLV or His986 in OmRV) (Fig. 5A). Therefore, these observations strongly suggest that the cap-snatching pocket no longer structurally exists in GLV and OmRV CPs and that the capsids of the *Totiviridae* viruses has lost the cap-snatching function in evolution. This finding is consistent with the lack of a 5’-cap RNA structure in GLV transcripts [15,33].

**Fig. 5.**
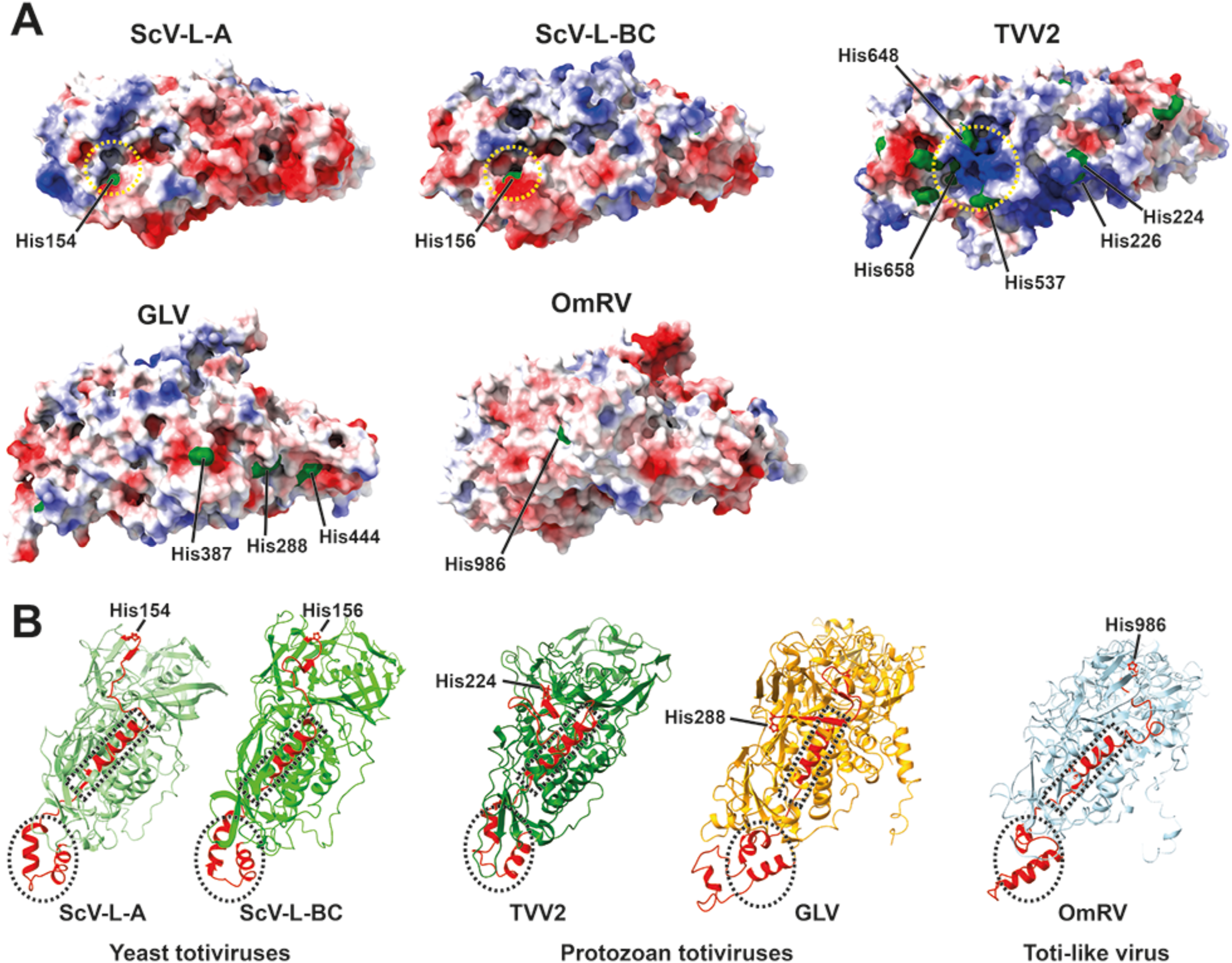
Lack of putative cap-snatching pockets in GLV and OmRV. **(A)** Electrostatic surfaces of *Totiviridae* and toti-like virus CPs are shown in blue (positive) and red (negative) scales. The CP of the yeast *Totiviridae* viruses ScV-L-A and ScV-L-BC have cap-snatching active pockets and invariant His residues (His154 or His156) (Yellow dotted circles). The CP of TVV2 is speculated to have a cap-snatching pocket with three putative His residues (His537, His648, and His658) in a similar position corresponding to the one in the yeast *Totiviridae* viruses (Yellow dotted circle). However, in GLV and OmRV CPs, no His residue is observed on the corresponding cap-snatching pocket, although three His residues (His288, His387, His444) in GLV CP and one His residue (His986) in OmRV CP are exhibited on the surfaces. (B) Conserved helices and invariant His residue in *Totivirdiae* and toti-like viruses. A conserved helix-turn helix and a long helix are highlighted as dotted circles and rectangles in each CP structure.

### Structural differences between GLV-HP and GLV-CAT

GLV-HP and GLV-CAT present remarkably distinct intracellular localization and extracellular release efficiency when chronically infecting the *G. duodenalis* isolate WBC6, which could, in turn, change their virulence [15]. The GLV-HP particles obviously interact with each other (Figs. 1B, 1I), which causes particle aggregation and affects the preferable intracellular localization [15]. Such particle-to-particle interactions should be mediated by the surface of the GLV-HP capsid; therefore, the structural differences between GLV-HP and GLV-CAT CPs are the center of the discussion.

Within the GLV-HP and GLV-CAT CP amino acid sequences, 45 amino acid alterations exist in the structurally visible region (Ile70–Val929) (Fig. 6A and Supplementary Fig. S6). Although the capsid surface of GLV-HP seems to be less charged (Fig. 4), these amino acid alterations do not dramatically alter the hydrophobicity and charges on the capsid surface. However, apparent structural differences are observed between GLV-HP and GLV-CAT CP-A/CP-B dimers (Fig. 6B, Supplementary Fig. S7). The major structural difference of CP-A is the aforementioned lack of C-terminal extension in GLV-HP (Fig. 2, Supplementary Fig. S7), while that of CP-B is a variable loop that consists in Pro316-Ser330 residues, localizing between the interfaces of CP-B and CP-A (Fig. 6B, Supplementary Fig. S7). Both CP-A and CP-B show high RMSD in the proximal to the 5-fold axis of the capsid (Fig. 6B, Supplementary Fig. S7).

**Figure 6.**
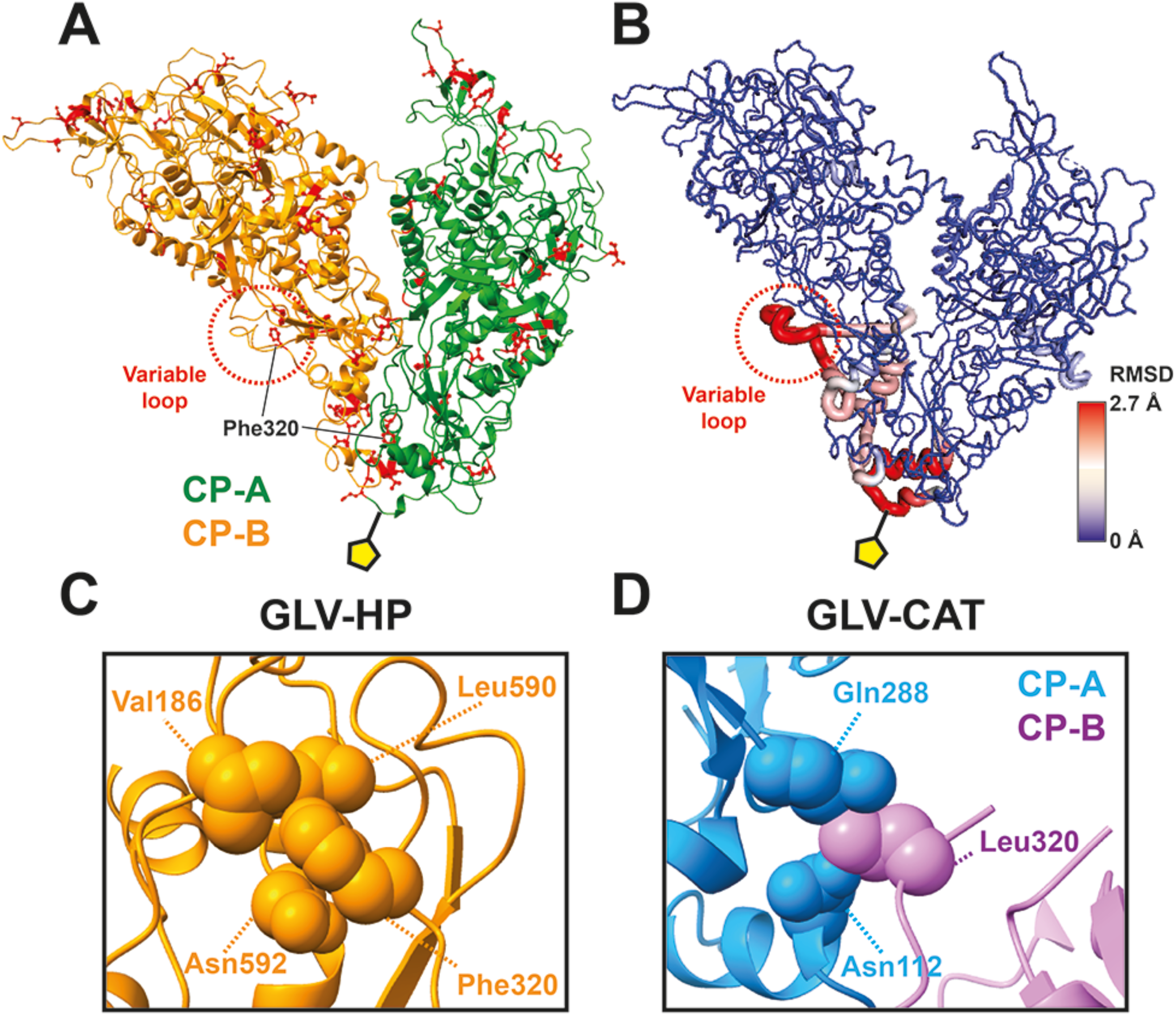
Amino acid alterations and structural variability between CPs in GLV-HP and GLV-CAT. (A) Amino acid alterations between the GLV-HP and GLV-CAT are mapped on a GLV-HP CP-A/CP-B dimer. (B) Local RMSD values are calculated per amino acid residue. The RMSD values are scaled by blue (low RMSD) and red (high RMSD). The yellow pentagon indicates the 5-fold axis of the GLV capsid. (C) Intra-subunit interaction of Phe230 of CP-B in the GLV-HP capsid. (D) Inter-subunit interaction between CP-A and Leu320 of CP-B in the GLV-CAT capsid.

Notably, Phe320 in GLV-HP forms intra-subunit interactions with Val184, Leu590, and Asn592 in the CP-B (Fig. 6C); however, in GLV-CAT, the corresponding Leu320 in CP-B interacts with Asn112 and Gln282 residues in the adjacent CP-A (Fig. 6D). The switching between intrasubunit and intersubunit interactions by the F320L point mutation largely contributes to driving the observed structural change in the variable loop. The Phe320 or Leu320 residue in the CP-A is located near the 5-fold axis, which might also affect the observed structural variation there (Figs. 6A-B). This additional CP–CP interaction that is mediated by Leu320 and the variable loop in GLV-CAT should make the CPs’ network in the capsid more robust. Together with the more robust C-terminal interlocking in the GLV-CAT capsid (Fig. 2), the capsid assembly of GLV-CAT is structurally more stable than that of GLV-HP. It is also speculated that lack of intersubunit interactions that are mediated by the variable loops within GLV-HP capsid might allow intersubunit interactions between two virions by them instead.

## Conclusion

The comparative analysis of the first high resolution atomic model of two GLV prototypes enables us to present structural insights into acquired and lost functional features in the evolution of dsRNA *Totiviridae* and toti-like viruses. The two lysine residues that compose the GLV pores are critical for understanding intraparticle genome synthesis and particle assembly in GLV and other *Totiviridae* viruses. The surface loops that exhibit only in the OmRV and the GLV capsid explain an essential requirement for acquiring extracellular cell-to-cell transmission, in particular virus entry and egress. The structural differences between the two GLV prototypes show significant conformational differences in the C-terminal extension and the variable loop that are critical for CP–CP interactions. It is hypothesized that the loose assembly of the CPs in GLV-HP could expose hydrophobic CP–CP interfaces to facilitate particle–particle interactions. This hypothesis is in agreement with observations in other viral model, such as the VLPs formed by VP40, the major capsid protein of (-)ssRNA Ebola virus, for which mutations at a specific loop in the protein N-terminus alters both protein oligomerization and virus egress [53]. It is interesting that these structural changes are potentially responsible for the transmission and virulence of the two GLV strains, which will provide strategies to engineer GLV to enhance its pathogenic effects on *G*. *duodenalis*.

## Materials and methods

### Parasite culture and viral particle preparation

The *G. duodenalis* isolates used in this study were HP-1 (Assemblage AI), originally isolated from a human patient [54], a kind gift of Prof. E. Noynkova (Charles University of Prague, Czechia), and CAT-1 (Assemblage AI), originally isolated from a cat [55], a kind gift of Dr. G.S. Visvesvara (Centers for Disease Control and Prevention, Atlanta, Georgia). HP-1 and CAT-1 are chronically infected with the virus strains GLV-HP and GLV-CAT, respectively [15]. Trophozoites were routinely grown in axenic, microaerophilic conditions in TYI-S33 medium, supplemented with 10% (v/v) adult bovine serum (Euroclone S.p.A., Milan, Italy) and 0.05% (v/v) bovine bile (Sigma-Aldrich, Merck Life Science S.r.l., Milan, Italy) at 37°C in 10 mL screw-cap tubes (NuncTM, Thermo Fisher Scientific, Waltham, MA, USA) and sub-cultured when confluence was reached (every 48–72 h). For viral particle purification, 1 L of trophozoite culture of each isolate was grown in 50 screw-cap tubes (Falcon, Thermo Fisher Scientific, Waltham, MA, USA) with TYI-S33 medium for 60 h. Tubes were chilled on ice to detach trophozoites, and the parasites were harvested by centrifugation at 900 × *g* for 10 min at 4°C. Trophozoite pellets were combined and washed twice with cold PBS by centrifugation at 900 × *g* for 10 min at 4°C. Trophozoite pellets were resuspended in 8 mL of cold PBS and lysed by sonication (5–6 times for 30 s at 60% power and 10% duty cycle) with a Sonoplus ultrasonic homogenizer (Bandelin Electronic, Berlin, Germany). The lysate was centrifuged at 10,400 × *g* (in JA20 centrifuge, Beckman) for 10 min at 4°C to remove debris. To sediment the viral particles, the supernatant was layered onto PBS/1.5 M sucrose solution in sample/sucrose 4:1 ratio and ultracentrifuged at 264,000 × *g* on Optima TLA 100.3 ultracentrifuge (Beckman for 2 h at 4°C. The virus pellet was resuspended with 8 mL of cold PBS, CsCl was added to a final density of 1.39 g/mL, and the volume was adjusted to 12 mL with PBS/CsCl solution at the same density. Virions were banded by density gradient centrifugation at 152,000 x *g* for 16 h at 4°C in a SW41 rotor (Beckman Coulter SRL, Milan, Italy). The gradient was then fractionated, and virion-positive fractions were identified by phenol extraction of nucleic acids, as previously described [15]. The positive fractions were pooled and dialyzed overnight in sterile PBS/glycerol 20% (v/v) and stored at -80°C until use. The overall quality of the prepared purified virions was examined by transmission electron microscopy (TEM), as previously described [15]. For the cryo-EM grid preparation, the purified GLV-HP and GLV-CAT particles were further concentrated down to 20–30 µL (approximately 10^9^– 10^10^ particles/mL) for the cryo-EM grid preparation.

### Cryo-EM data acquisition and map reconstruction

Three microliters of the purified GLV-HP or GLV-CAT sample was placed on a glow-discharged holey carbon grid (Quantifoil R2/2 or R2/1, Cu 300 mesh; Quantifoil Micro Tools GmbH) for 3 s blotting time at 4°C, 100% humidity with filter paper, and then plunge-freezing in liquid ethane using Vitrobot Mark IV (Thermo Fisher Scientific). The prepared grids were pre-screened with a 200-kV Glacios cryo-EM at the Uppsala cryo-EM center. The full datasets of GLV-HP and GLV-CAT were collected with a 300-kV Titan Krios G2 cryo-EM (Thermo Fisher Scientific) equipped with a Gatan K3 BioQuantum detector and the post-column electron energy filter (20 eV slit width) at the SciLifeLab Cryo-EM infrastructure unit. The image movies were collected by a counted super-resolution mode at a nominal magnification of 81,000×, which corresponds to a pixel size of 1.06 Å /pixel. The defocus range is 0.2 µm steps in the range of 0.7–1.5 µm under focus. The total exposure per movie was adjusted to 30 e^-^/Å^2^ for 1.9 s and dose-fractionated into 30 frames. A total of 19,580 and 7,098 movies were collected for further image analysis, respectively. The parameters of the cryo-EM data collection are shown in Supplementary Table S1. The three-dimensional (3D) cryo-EM map of the capsid was reconstructed using CryoSPARC version 4.3.1 [56] linked to a local GPU/CPU computer cluster. The image frames were corrected by the patch motion correction using frames 3-28, and then the contrast transfer function (CTF) parameters were estimated using the patch CTF. The first template images of the virus particles were generated from 100–200 manually picked particles, and the templates were used for automatically picking up virus particles using the template picker. Both the picked GLV-HP and the GLV-CAT images contain approximately 20% empty particles; otherwise, all of them are filled particles. After a couple of 2D classifications, good 2D classes containing 28,342 and 5,804 good particles of GLV-HP and GLV-CAT, respectively, were selected. The final 3D map reconstruction was accomplished using the Homogeneous Refinement option with imposing icosahedral symmetry and an initial model that was generated by 20-Å lowpass filtering to a previously determined OmRV cryo-EM map [25]. Throughout the reconstruction process, the per-particle defocus, CTF parameters, spherical aberration, beam tetrafoil, and beam anisotropic magnification were optimized to enhance map qualities. The final 3D maps of GLV-HP and GLV-CAT were determined to have a resolution of 2.1 Å and 2.6 Å, respectively, using the gold standard Fourier shell correlation (FSC) at a 0.143 cutoff (Supplementary Fig. S1). These maps were subsequently employed to construct atomic models of the CPs.

### Atomic modeling and refinement

The first atomic models of GLV CPs (A and B subunits) were manually built in the cryo-EM map of GLV-HP using Coot version 1.0.06 [57]. The manually built atomic model was further refined iteratively using PHENIX 1.20.1 [58] and Coot. The amino acid residues of the refined atomic model of GLV-HP CPs were then mutated to those of GLV-CAT and placed on a GLV-CAT cryo-EM map. The GLV-CAT CPs were then refined using PHENIX and COOT, as described above. The validation statistics of the atomic models and the cross-correlation to the cryo-EM maps are shown in Supplementary Table S1.

### Structural analysis

To render the cryo-EM models and atomic models, the UCSF Chimera and ChimeraX were used [59,60]. The viral CP structures resembling that of GLV-HP were comprehensively searched against all PDB-deposited structures using the Dali structure comparison server [61]. Eight CP structures in non-enveloped icosahedral dsRNA viruses were detected to be similar to the CP of GLV-HP. These eight CP structures and the GLV-HP structure were superimposed by pairwise structure-based alignments using the MUSTANG program [62]. During the alignments, all-to-all RMSD values were calculated in the superimposed structures. These RMSD values were used as a distant matrix to generate a structural phylogeny of the viral CPs using a neighbor-joining method in MEGA X [63], as already described earlier [64,65]. The aligned structures were also visualized to discover conserved and acquired unique structures in the GLV CP. To calculate the local RMSD, the local_rms command was applied for the aligned GLV-HP and GLV-CAT CP dimers using the PyMOL script collection (PSICO) module [66].

## Data availability

The cryo-EM maps of the GLV-HP and the GLV-CAT are available in the EMDB database, entries EMD-18791 and EMD-18792. The atomic models of the CPs are available in the PDB database, entries 8R0F and 8R0G.

## Acknowledgments

The data was collected at the Cryo-EM Swedish National Facility funded by the Knut and Alice Wallenberg, Erling Persson Family, and Kempe Foundations, SciLifeLab, Stockholm University and Umeå University. We thank Julian Conrad for help with data acquisition. We thank Björn Persson for reconstructing the initial low-resolution cryo-EM maps of the GLV-HP and the GLV-CA; Prof. Eva Nohýnková, Charles University of Prague, Czechia, and Dr. Govinda S. Visvesvara, Centers for Disease Control and Prevention, Atlanta, Georgia, for kindly gift of HT-1 and CAT-1 G. duodenalis isolates, respectively. A.M. acknowledges support from DESY (Hamburg, Germany), a member of the Helmholtz Association HGF.

## Financial enclosures

Funding was provided by following agencies: the Swedish Research Council (to K.O., grant number: 2018-03387; to A.M., grant number: 2022-00236), the Swedish Foundation for International Cooperation in Research and Higher Education (STINT) (to Janos Hajdu and K.O., grant number: JA2014-5721), FORMAS research grant from the Swedish Research Council for Environment, Agricultural Sciences and Spatial Planning (to K.O., grant number: 2018-00421), Royal Swedish Academy of Sciences (to K.O., grant number: BS2018-0053), Norwegian Research Council (to K.O., grant number: 324266), Istituto Superiore di Sanità (to M.L., grant number: ISS20-4389733b36a1).

## Author contributions

M.G. and M.L. prepared samples. H.W., A.M., M.L. and K.O. wrote the manuscript. M.L. and K.O. designed the experiments. H.W., A.M. and K.O. collected the cryo-EM data and H.W., M.M.H. and K.O. analyzed the data. All of the authors discussed the results and proofread the manuscript.

## Supporting information captions

**Supplementary Fig. S1. Overall and local resolution estimations of GLV-HP and GLV-CAT reconstructions. (A)** FCS curves and (B) local resolution of the final cryo-EM 3D reconstruction for GLV-HP and GLV-CAT.

**Supplementary Fig. S2. Assigned amino acid residues and the secondary structure diagram of the atomic model of the CP-B of GLV-HP and GLV-CAT.**

**Supplementary Fig. S3. Structural comparison of CP-A and CP-B.** The CP-A and CP-B of GLV-HP are colored in green and orange, and those of GLV-CAT are colored in light blue and purple. The total RMSD values between CP-A and CP-B were calculated for GLV-HP and GLV-CAT. Some conformational changes are observed in the regions indicated by red dotted circles.

**Supplementary Fig. S4 DALI search and RMSD-based structural phylogeny of CPs in non-enveloped icosahedral dsRNA viruses. (A)** A list of the eight identified CPs that are similar to the GLV CP obtained from a Dali search (Z score: 3.4–10.4). (B) RMSD-based structural phylogeny generated by the alignment of the GLV CP with the eight CP structures in a Dali search.

**Supplementary Fig. S5 Selected 2D class-averaged projections of capsid-subtracted GLV-HP particle images.** The striped genome (side views, red squares as examples) and the spiral genome (top views, blue squares as examples) are observed, which are typical 2D classes of partially spooled genome organization in Reoviridae viruses.

**Supplementary Fig. S6 Amino acid sequence alignment of capsid proteins between the GLV-HP and GLV-CAT strains.** The alignment was generated by the CLC sequence viewer 7.

**Supplementary Fig. S7 Structural differences of CP-A and CP-B in GLV-HP and GLV-CAT.** The overall RMSD was calculated to evaluate the structural similarity. Conformational changes are observed in the regions (C-terminal extension in CP-A and variable loop in CP-B), indicated by yellow dotted circles.

**Supplementary Table S1. Cryo-EM data collection, refinement, and validation statistics.**

