## Supplementary information (captions, figures, table) for "High-resolution comparative atomic structures of two Giardiavirus prototypes infecting *G. duodenalis* parasite"

**Supporting information captions**

**Supplementary Fig. S1. Overall and local resolution estimations of GLV-HP and GLV-CAT reconstructions. (A)** FCS curves and (B) local resolution of the final cryo-EM 3D reconstruction for GLV-HP and GLV-CAT.

**Supplementary Fig. S2. Assigned amino acid residues and the secondary structure diagram of the atomic model of the CP-B of GLV-HP and GLV-CAT.**

**Supplementary Fig. S3. Structural comparison of CP-A and CP-B.** The CP-A and CP-B of GLV-HP are colored in green and orange, and those of GLV-CAT are colored in light blue and purple. The total RMSD values between CP-A and CP-B were calculated for GLV-HP and GLV-CAT. Some conformational changes are observed in the regions indicated by red dotted circles.

**Supplementary Fig. S4 DALI search and RMSD-based structural phylogeny of CPs in non-enveloped icosahedral dsRNA viruses. (A)** A list of the eight identified CPs that are similar to the GLV CP obtained from a Dali search (Z score: 3.4–10.4). (B) RMSD-based structural phylogeny generated by the alignment of the GLV CP with the eight CP structures in a Dali search.

**Supplementary Fig. S5 Selected 2D class-averaged projections of capsid-subtracted GLV-HP particle images.** The striped genome (side views, red squares as examples) and the spiral genome (top views, blue squares as examples) are observed, which are typical 2D classes of partially spooled genome organization in Reoviridae viruses.

**Supplementary Fig. S6 Amino acid sequence alignment of capsid proteins between the GLV-HP and GLV-CAT strains.** The alignment was generated by the CLC sequence viewer 7.

**Supplementary Fig. S7 Structural differences of CP-A and CP-B in GLV-HP and GLV-CAT.** The overall RMSD was calculated to evaluate the structural similarity. Conformational changes are observed in the regions (C-terminal extension in CP-A and variable loop in CP-B), indicated by yellow dotted circles.

**Supplementary Table S1 Cryo-EM data collection, refinement, and validation statistics.**

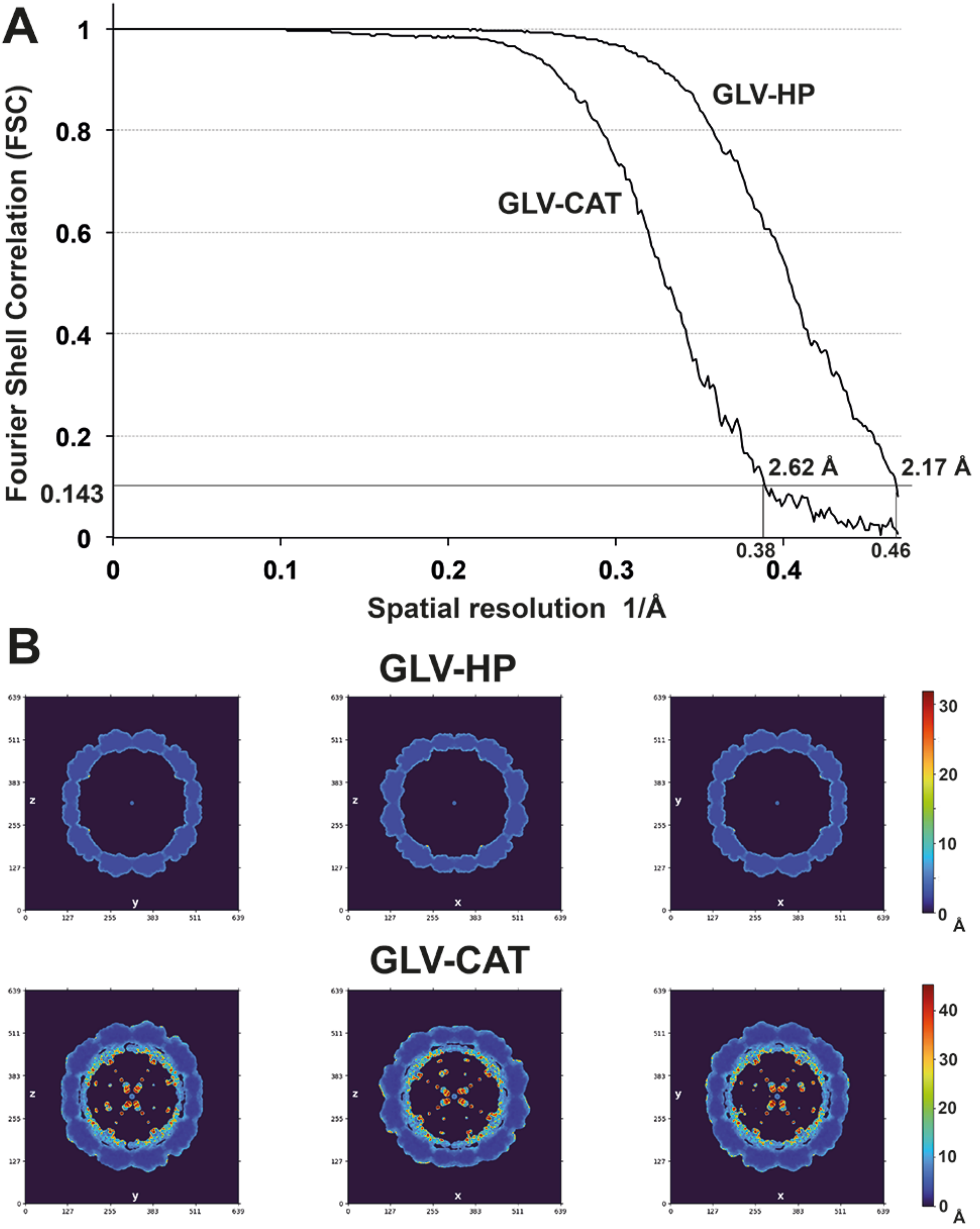
A
Fourier Shell Correlation (FSC)
1
0.8
0.6
0.4
0.2
0.143
0
0
0.1
0.2
0.3
0.4
0.38
0.46
Spatial resolution 1/Å
GLV-HP
GLV-CAT
2.62 Å
2.17 Å
B
GLV-HP
GLV-CAT
0
10
20
30
40
Å

Fig. S1

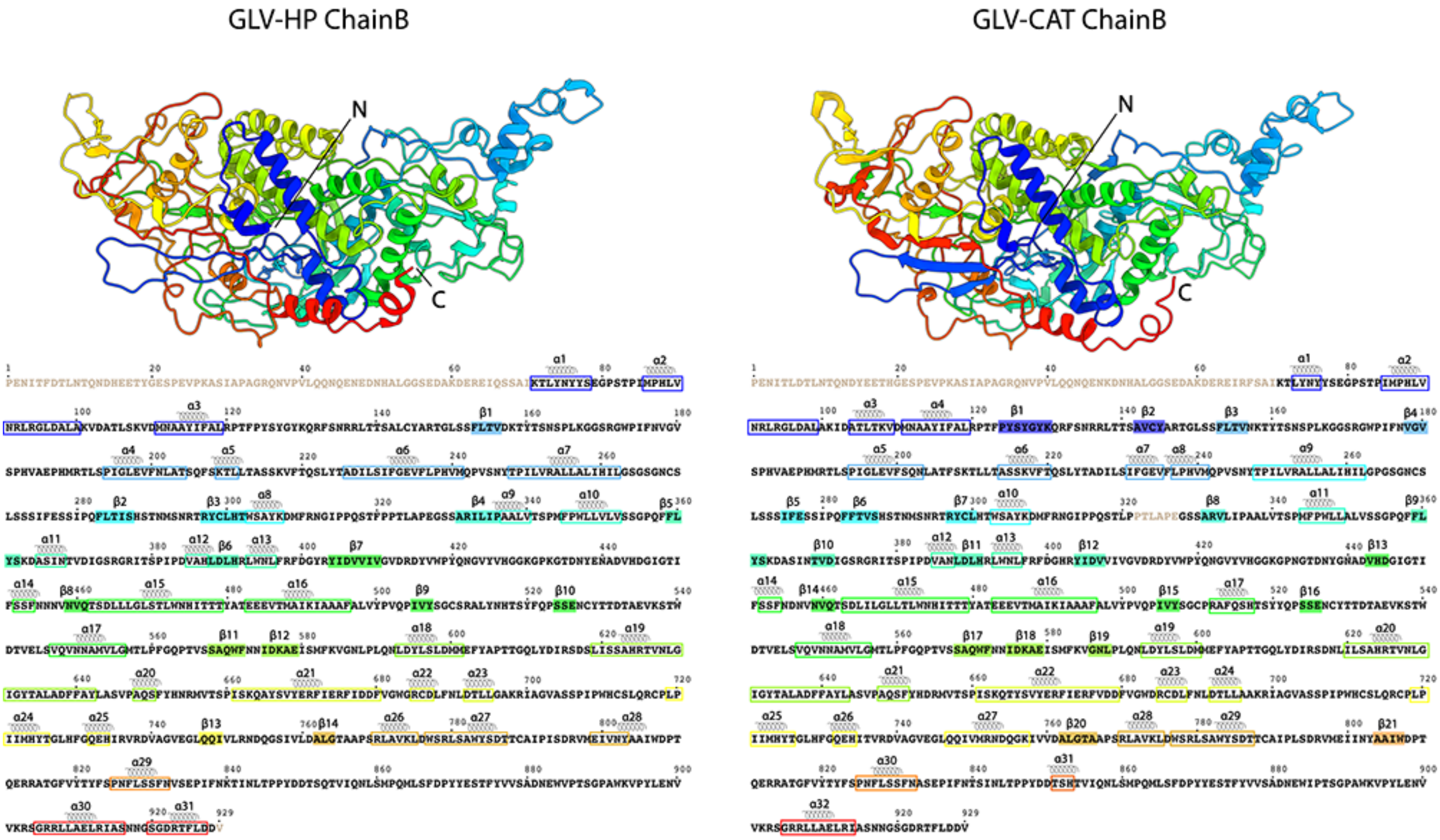


Fig. S2

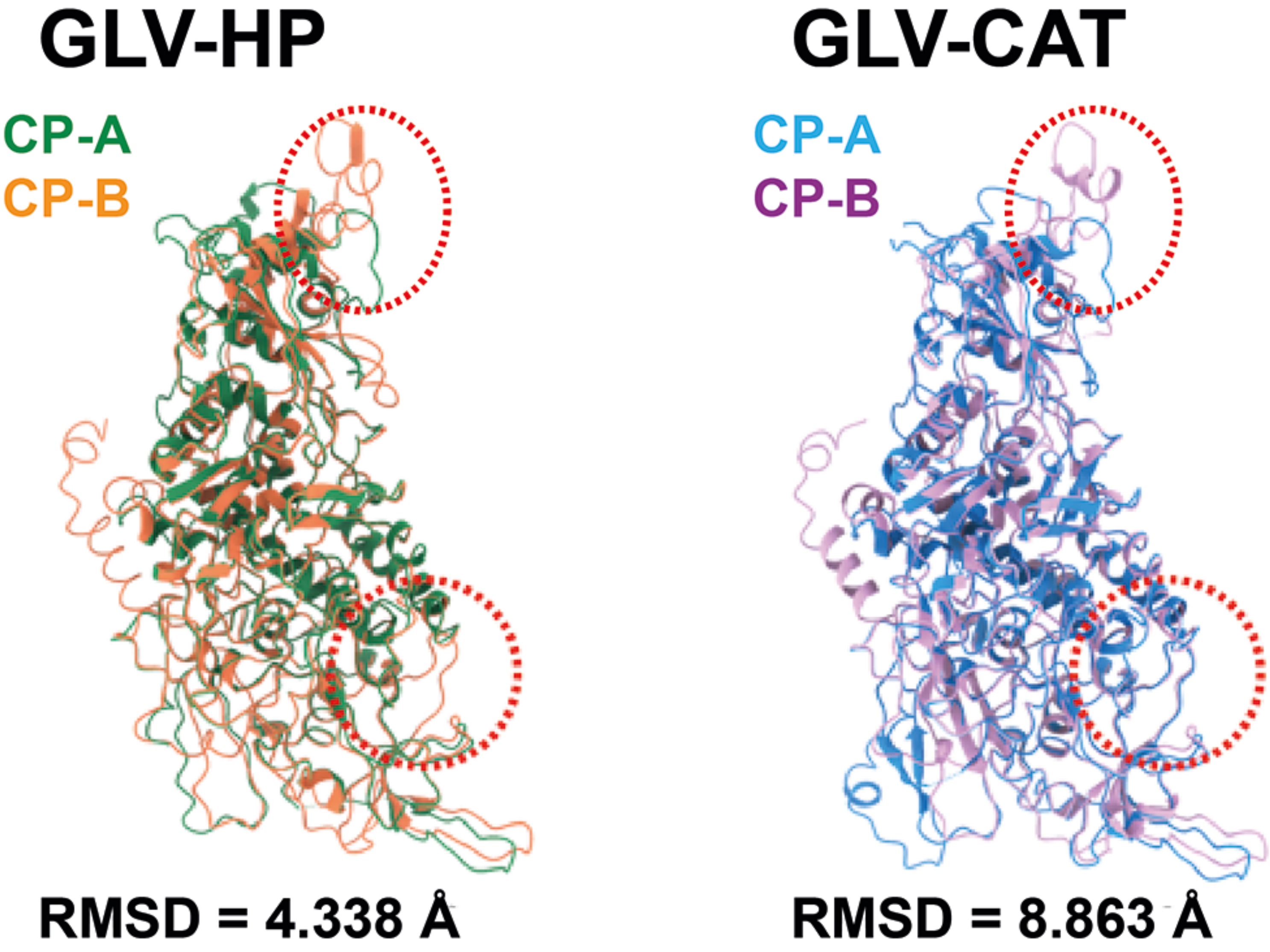


Fig. S3

**A**

| Virus species | Taxon | Z score | PDB ID |
|---|---|---|---|
| Omono River Virus (OmRV) | Toti-like virus (Unclassified Totiviridae) | 10.4 | 7d01 |
| Penicillium Chrysogenum virus (PcV) | Chrysoviridae | 7.1 | 3j3i |
| Saccharomyces cerevisiae virus L-A (ScV-L-A) | Totiviridae | 6.3 | 1m1c |
| Saccharomyces cerevisiae virus L-BC (ScV-L-BC) | Totiviridae | 5.8 | 7qwx |
| Leishmania RNA virus 1 (LRV1) | Totiviridae | 4.5 | 7ns2 |
| Rosellinia necatrix megabirnavirus 1 (RnMBV1) | Megabirnaviridae | 4.5 | 8b4z |
| Trichomonas Vaginalis Double-Stranded RNA Virus | Totiviridae | 4.4 | 7lwy |
| Rosellinia necatrix quadrivirus 1 (RnQV1) | Quadriviridae | 3.4 | 5nd1 |

**B**

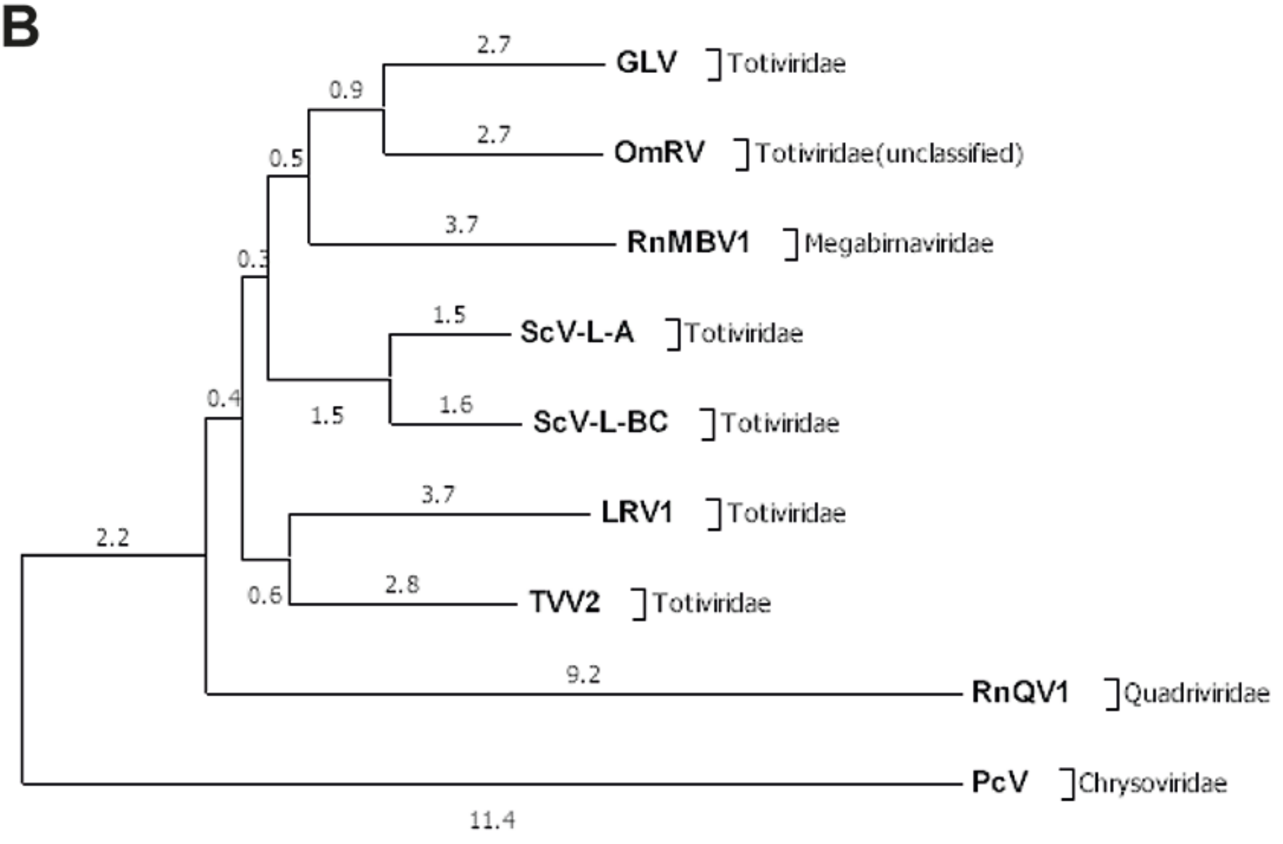


**Fig. S4**

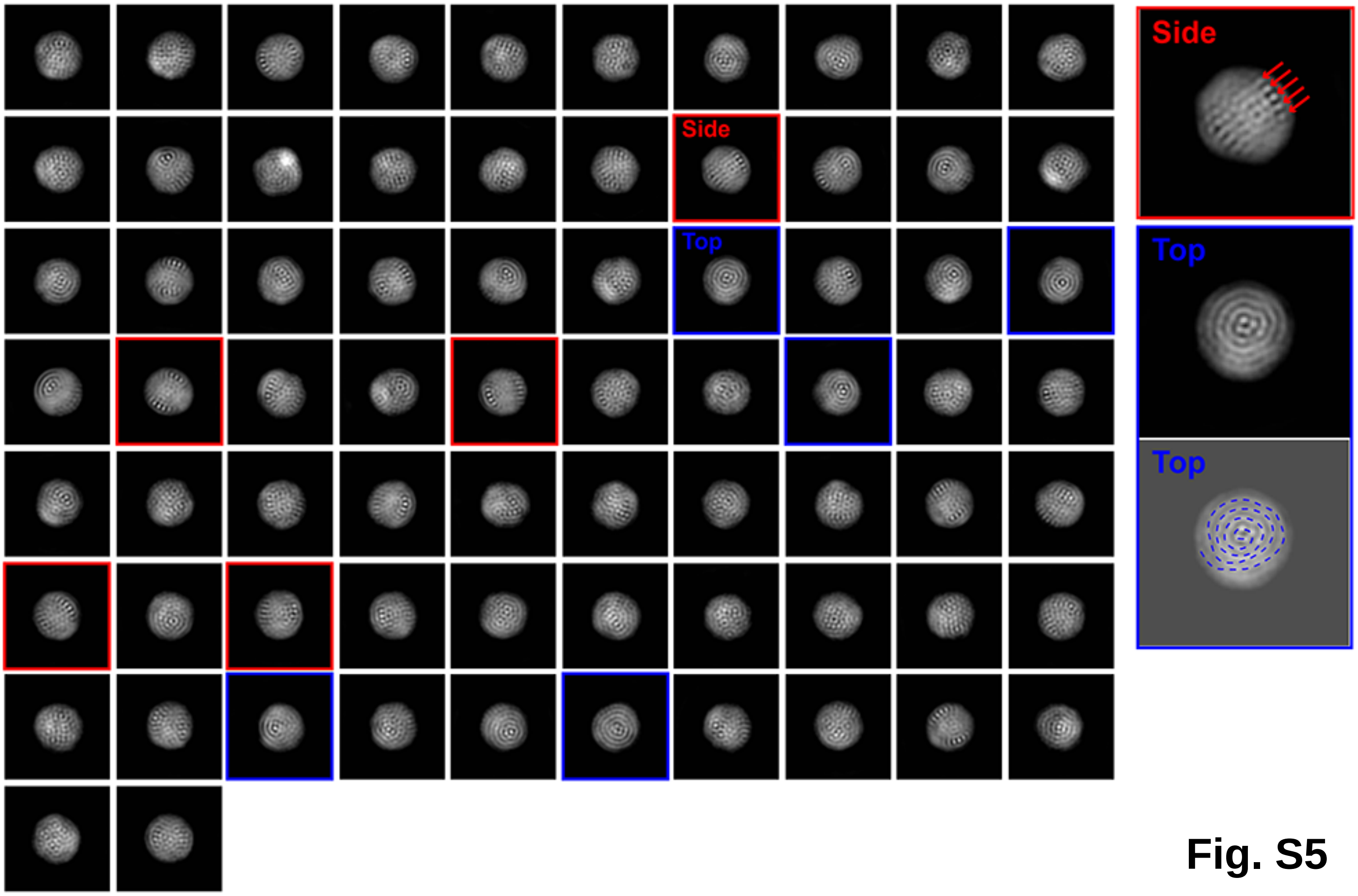


Fig. S5

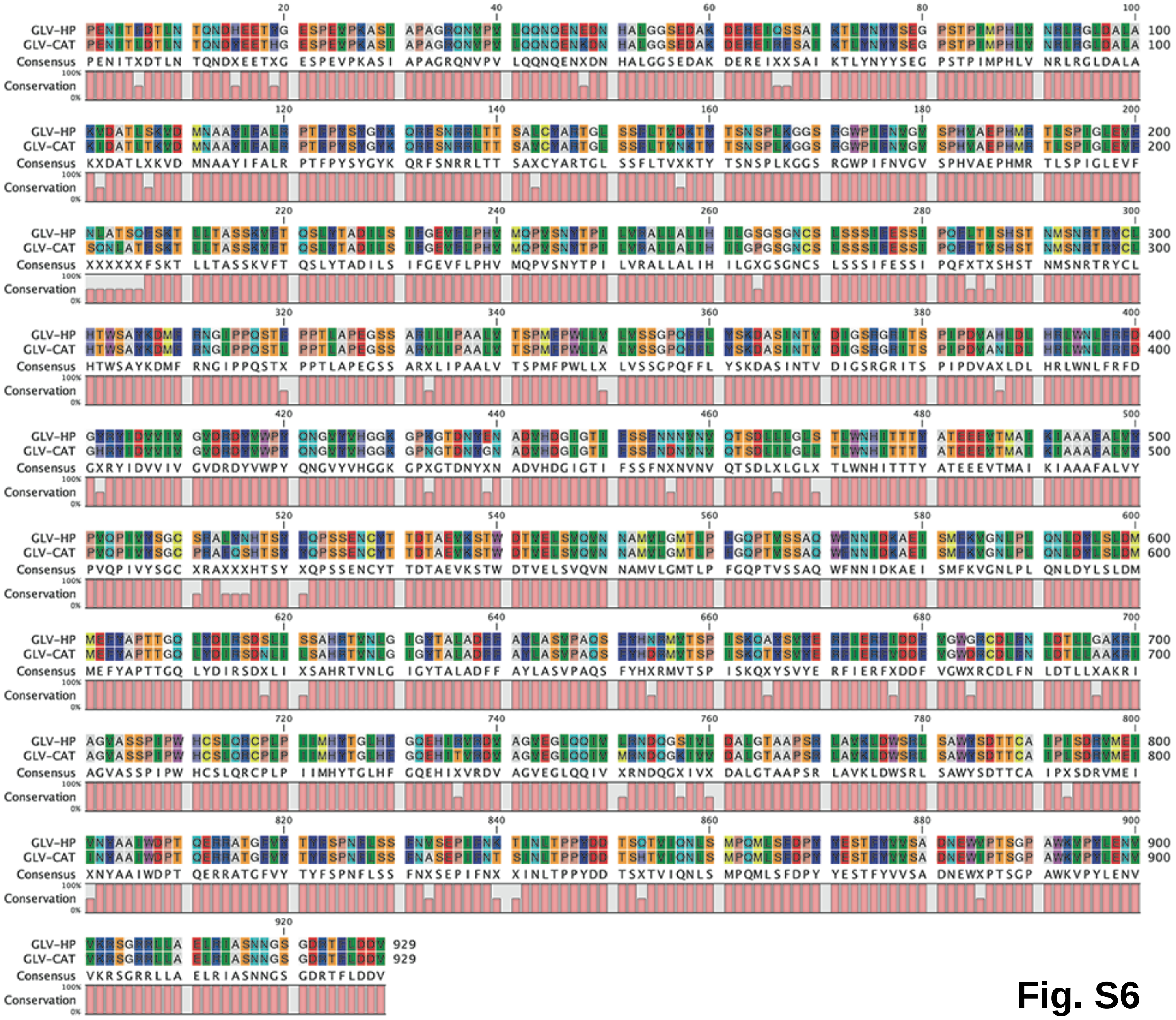
GLV-HP
GLV-CAT
Consensus
Conservation
100%
0%
20 40 60 80 100
PENITXDTLN TQNDXEETXG ESPEVPKASI APAGRQNVPV LQQNQENXDN HALGGSEDAK DEREIXXSAI KTLYNYYSEG PSTPIMPHLV NRLRGLDALA
120 140 160 180 200
KXDATLXKVD MNAAYIFALR PTFPYSYGYK QRFSNRRLTT SAXCYARTGL SSFLTVXKTY TSNSPLKGGS RGWPIFNVGV SPHVAEPHMR TLSPIGLEVF
220 240 260 280 300
XXXXXXFSKT LLTASSKVFT QSLYTADILS IFGEVFLPHV MQPVSNYTPI LVRALLALIH ILGXGSGNCS LSSSIFESSI PQFXTXSHST NMSNRTRYCL
320 340 360 380 400
HTWSAYKDMF RNGIPPQSTX PPTLAPEGSS ARXLIPAALV TSPMFPWLLX LVSSGPQFFL YSKDASINTV DIGSRGRITS PIPDVAXLDL HRLWNLFRFD
420 440 460 480 500
GXRYIDVVIV GVDRDYVWPY QNGVYVHGGK GPXGTDNYXN ADVHDGIGTI FSSFNXNVNV QTSDLXLGLX TLWNHITTTY ATEEEVTMAI KIAAAFALVY
520 540 560 580 600
PVQPIVYSGC XRAXXXHTSY XQPSSENCYT TDTAEVKSTW DTVELSVQVN NAMVLGMTLP FGQPTVSSAQ WFNNIDKAEI SMFKVGNLPL QNLDYLSLDM
620 640 660 680 700
MEFYAPTTGQ LYDIRSDXLI XSAHRTVNLG IGYTALADFF AYLASVPAQS FYHXRMVTSP ISKQXYSVYE RFIERFXDDF VGWXRCDLFN LDTLLXAKRI
720 740 760 780 800
AGVASSPIPW HCSLQRCPLP IIMHYTGLHF GQEHIXVRDV AGVEGLQQIV XRNDQGXIVX DALGTAAPSR LAVKLDWSRL SAWYSDTTCA IPXSDRVMEI
820 840 860 880 900
XNYAAIWDPT QERRATGFVY TYFSPNFLSS FNXSEPIFNX XINLTPPYDD TSXTVIQNLS MPQMLSFDPY YESTFYVVSA DNEWXPTSGP AWKVPYLENV
920
VKRSGRRLLA ELRIASNNGS GDRTFLDDV 929

Fig. S6

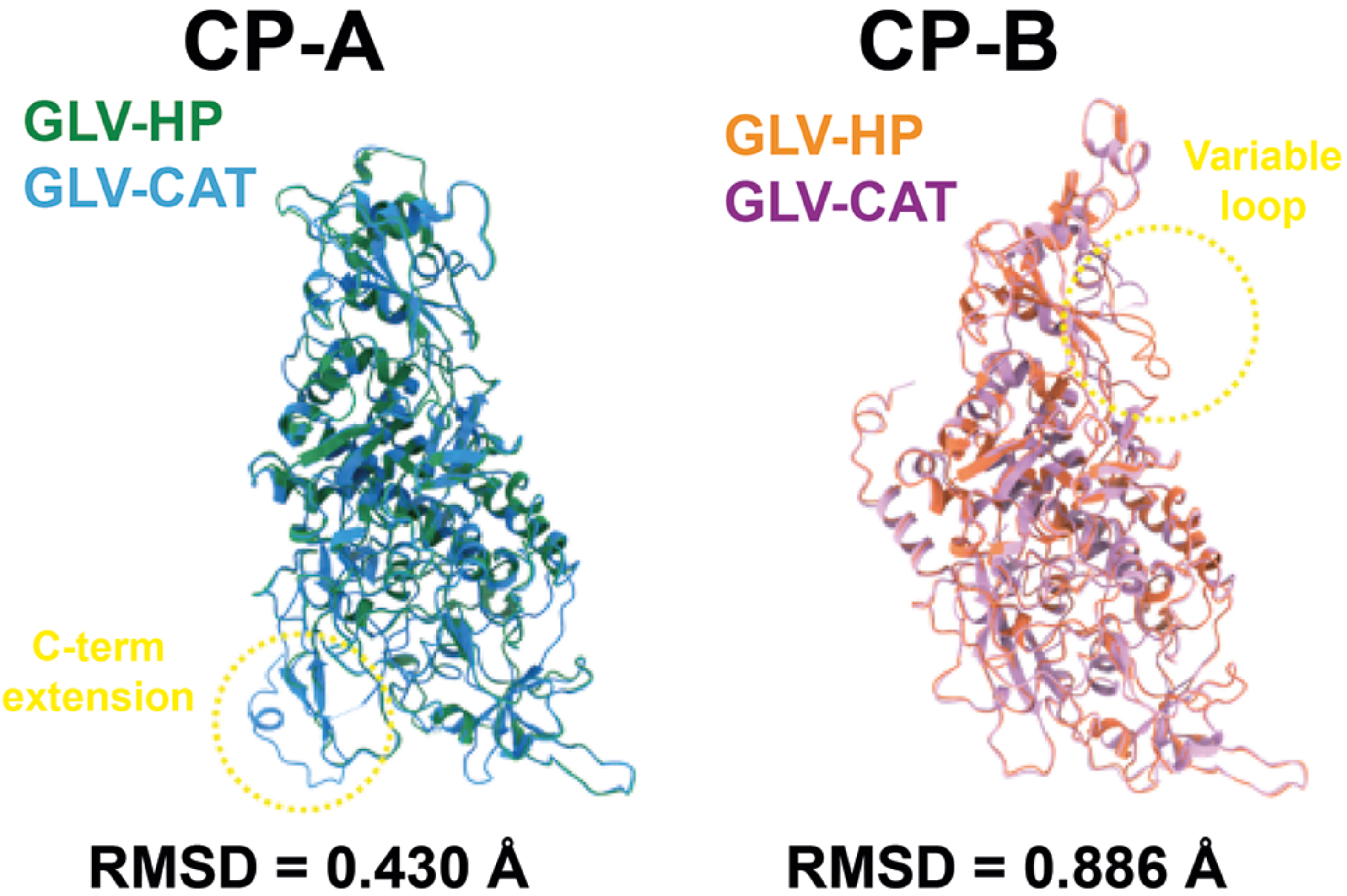
CP-A
GLV-HP
GLV-CAT
C-term
extension
RMSD = 0.430 Å
CP-B
GLV-HP
GLV-CAT
Variable
loop
RMSD = 0.886 Å

Fig. S7

**Table 1. Cryo-EM data collection, refinement, and validation statistics.**

| Data collection and processing | GLV-HP | GLV-CAT |
|---|---|---|
| Magnification | 81,000 | 81,000 |
| Voltage (kV) | 300 | 300 |
| Electron exposure ($e^{-}$/Å$^2$) | 30 | 30 |
| Defocus range (μm) | -0.7 to -1.5 | -0.7 to -1.5 |
| Pixel size (Å) | 1.06 | 1.06 |
| Symmetry | I | I |
| Final particle (No.) | 28,342 | 5,804 |
| Map resolution (Å) | 2.14 | 2.62 |
| FSC threshold | 0.143 | 0.143 |
| Model composition and quality | | |
| Non-hydrogen atoms | 12,852 | 13,445 |
| Protein residues | 1,633 | 1,708 |
| B-factors (Å$^2$) | | |
| Protein (min/max/mean) | 0/44/11.07 | 8.05/98.34/42.21 |
| RMSD* | | |
| Bond lengths (Å) | 0.011 | 0.005 |
| Bond angles (°) | 1.112 | 0.655 |
| Validation | | |
| MolProbity score | 2.21 | 2.59 |
| Clashscore | 6.68 | 8.88 |
| Poor rotamers (%) | 3.64 | 9.18 |
| Ramachandran plot | | |
| Favored (%) | 93.73 | 94.35 |
| Allowed (%) | 5.41 | 4.77 |
| Outliers (%) | 0.86 | 0.88 |
| CC (mask) | 0.83 | 0.85 |
| EMRinger score | 4.32 | 3.39 |
| Data deposition | | |
| EMDB | EMD-18791 | EMD-18792 |
| PDB | 8R0F | 8R0G |

*RMSD, root mean square deviation

**Table S1**
